# Is facultative sex is the best of both worlds in the parasitoid wasp *Lysiphlebus fabarum?*

**DOI:** 10.1101/2024.02.25.581911

**Authors:** Rebecca A Boulton

**Affiliations:** Biological and Environmental Sciences, University of Stirling, FK9 4NF, United Kingdom; Laboratory of Genetics, Plant Sciences Group, Wageningen University and Research, 6708 8 PB, Netherlands

## Abstract

The prevalence of sexual reproduction has long puzzled evolutionary biologists. This is because asexual reproduction is a far more efficient way to transmit copies of one’s own genome to the next generation, and, all-else-being-equal, should predominate over sex. Asexual reproduction is not without its disadvantages though, the lack of genetic recombination can render parthenogenetic lineages vulnerable to extinction under environmental change, or compromise fitness due to the buildup of deleterious recessive mutations. Facultative sex, where individuals retain the ability to reproduce sexually and asexually, has been touted as ‘the best of both worlds’, providing the long-term genetic benefits of sex without the short-term costs. In this study I tested whether sex is facultative in the aphid parasitoid *Lysiphlebus fabarum*, and if so whether facultative sex is ‘the best of both worlds’. I found that females from asexual populations mate with males produced by females from an obligately sexual population, and produced offspring with paternal genetic contributions, confirming facultative sex in this species. I then assayed the fitness of these facultatively sexual wasps and found, contrary to predictions that they actually had significantly lower fitness than asexual or obligately sexual females, as their daughters were up to 40% more likely to fail to reproduce entirely. In contrast, obligately sexual females were not at a disadvantage compared to asexual females in this species; they had increased fecundity compared to asexual females which compensated for the costs of sex. In *L. fabarum* it seems that all is not equal; the fecundity advantage that obligate sex provides, along with the costs of occasional facultative sex by asexuals may balance the scales and allow stable coexistence of the different reproductive modes in nature.

## 1. Introduction

Understanding how sex became the dominant reproductive strategy in eukaryotes has long been a major theme in evolutionary biology [1–3]. The initial interest in this question arose from the observation that asexual reproduction should, in theory, predominate because it is a far more efficient way to transmit copies of one’s own genome to the next generation. An asexually reproducing individual is expected to have double the genetic representation of itself in the next generation compared to an equivalent sexual individual. This became known as the ‘twofold cost of sex’ [1]; early explanations considered ‘genome dilution’ to be the cause of this cost, whereby a sexually reproducing female ‘dilutes’ her genetic contribution to the next generation, halving her fitness compared to an asexual female. It is now accepted that ‘genome dilution’ is not the source of the ‘twofold’ cost of sex (as only genes for reproduction matter in this context, see [4] for an in-depth explanation), instead the twofold cost arises because sexual individuals must ‘waste’ resources on producing males, which do not efficiently convert resources to offspring [4]. When the sex ratio is 50:50, sexuals produce half as many daughters as asexuals, so the twofold cost of sex is sometimes known as the ‘twofold cost of males’ [1–4].

Given this cost, how then did sex become more prevalent than asexual reproduction in eukaryotes after it evolved over two billion years ago? And more recently, given that asexual reproduction readily re-evolves in eukaryotes [2] why does sex still dominate? Historically, two broad models which encompass different types of benefits have been proposed to answer these questions. The first group are the mutational models, which emphasize the importance of sex in exposing mutations to selection, so that deleterious mutations can be purged, and beneficial mutations can be fixed more effectively [3,5–8]. The second group are ecological models, which emphasize the benefits of sex for adaptive potential; by creating new gene combinations sex facilitates adaptation under environmental change [9]. Mutational and ecological benefits of sex are not mutually exclusive however and can interact in complex and context-dependent ways, which may explain why phylogenetic patterns point to reproductive mode being more evolutionarily labile than expected in the eukaryotes [2–4,10,11].

The picture becomes more complex in light of mounting evidence that many species thought only to reproduce through obligate sex or asexual reproduction are actually sexually plastic [10]. Many species reproduce asexually only some of the time, either as an obligate part of their lifecycle (as in cyclical parthenogens like many aphids and *Daphnia*; [11]), facultatively when a suitable mate can’t be found [12], or sometimes spontaneously (so-called *Tychoparthenogenesis* [2,13]. Facultative sex has been touted as the ‘best of both worlds’ as it can, in theory, provide both the short-term demographic benefits of asexual reproduction and the long-term genetic benefits of sex [11]. Integrating sexual plasticity in all its forms into theoretical and phylogenetic models which aim to reconstruct the evolution of reproductive mode is the obvious next step in answering the longstanding question ‘why sex?’, but this requires more data, not only about the occurrence of facultative sex (which is often cryptic), but also its costs and benefits relative to obligate strategies [4].

The sexual-asexual aphid parasitoid wasp *Lysiphlebus fabarum* offers an ideal system to explore whether facultative sex really does represent ‘the best of both worlds’. We already have a comprehensive understanding of the molecular basis and population genetics of reproductive mode in *L. fabarum* [14–16]. Asexual reproduction seems to have initially arisen 0.5 million years ago from ancestral obligate sex in *L. fabarum*, and it is determined by the inheritance of a single recessive allele which results in parthenogenesis by central-fusion automixis (the haploid products of meiosis I fuse to form diploid gametes which develop without fertilisation; [16,17]). Whilst this form of parthenogenesis does maintain heterozygosity to some extent, because recombination still occurs, asexual lineages can only become more homozygous, eventually resulting in inbreeding depression and associated reductions in fitness [2]. However, population genetic studies show that parthenogenetic populations of *L. fabarum* have high levels of heterozygosity and allelic variation even compared to sexuals [14,15]. This genetic diversity has been proposed to exist due to occasional bouts of rare sex by normally asexual females, as well as ‘contagious’ parthenogenesis, where rare males produced by asexual females cross with sexual females, creating new asexual lineages [14,15]. Regardless of the mechanism, this genetic variation is thought to underlie the success of asexual reproduction in this species, resulting in parthenogenetic populations dominating across Europe [14,15]. Obligately sexual populations are less commonly encountered in the field, but sex does persist in *L. fabarum*, and even occurs in sympatry with asexuality in some regions [17, 18].

This study aims to test whether facultative sex occurs in *L. fabarum* and if it does, whether it represents the ‘best of both worlds’, providing the short-term benefits of asexual reproduction and long-term genetic benefits associated with sex [12]. To do this I gave female *L. fabarum* from 7 asexual lines and one outbred sexual population aphid hosts to parasitise after (i) being exposed to males (from the sexual population) or (ii) being kept in isolation (due to haplodiploidy sexual females can produce unfertilised eggs that develop as male offspring). I found that asexual females copulated when they were exposed to males and so in a follow-up experiment, I genotyped their G1 offspring to test whether mating reflects a behavioural vestige of sexual reproduction without fertilisation or whether these females can still reproduce sexually. I also assayed the reproductive success of daughters produced by asexual females which either mated or remained virgin. This allowed me to test whether putative facultative sex allows females to reap the short-term demographic benefits of asexual reproduction (not producing males) and also gain the long-term genetic benefits of sex (which only appear in hybrid offspring).

## 2. Methods

### (a) Study system

In the aphid parasitoid *Lysiphlebus fabarum* both sexual and asexual populations exist, separately as well as in sympatry in some regions [14,15]. Asexual reproduction is inherited via a single recessive allele which appeared approximately 0.5 million years ago [14]. Inheritance of two copies of the asexual allele results in parthenogenetic asexual reproduction through central-fusion automixis, where the products of meiosis 1 fuse and recombine, restoring diploidy without integrating paternal DNA [16]. A panel of microsatellite markers has been developed to study the population genetics of *L. fabarum*, one of which, *Lysi07*, shows perfect linkage to the recessive genetic factor associated with asexual reproduction [18].

### (b) Lab rearing

This study used females from 7 asexual lines and males and females from an outbred obligately sexual population of the aphid parasitoid *Lysiphlebus fabarum*. These lines were provided by Prof Christoph Vorburger (EAWAG, Zürich, Switzerland) who has been maintaining them since they were collected between 2006 and 2017 (see [16] for details). The wasps have been cultured at the University of Stirling under equivalent conditions since March 2023 and are regularly genotyped to check their genetic identity has been maintained. The wasps are given black bean aphid (*Aphis fabae*) colonies to parasitise; they are able to parasitise all instars of aphid but prefer 1^st^ instar nymphs [19]. Female wasps lay a single egg inside each aphid (they are solitary endoparasitoids) and the host continues to grow as the wasp larva develops (they are koinobionts; [19]). In the lab aphids are maintained on broad bean seedlings (*Vicia faba*). All insects are kept in climate-controlled chambers at 20°C, 70% RH on a 16:8 hour light:dark cycle. Under these conditions the generation time of *L. fabarum* is 12-14 days.

### (c) Experimental design

To measure the benefits and costs of sex in *L. fabarum* I assayed offspring production by obligately sexual and asexual *L. fabarum* females (referred to as G0) which were exposed to a male (from the obligately sexual population), M, or remained virgin, V (fig 1a). All females (including asexuals) in the M treatment readily copulated with the male, so I refer to these females as ‘mated’. To test for longer-term costs of mating (and potentially facultative sex) I repeated the same fitness assays with the unmated daughters of V and M asexual females (fig 1c; V_2_ and M_2_; referred to as G1 females). The fitness measures I used were: (i) the number of aphid mummies (parasitised dead aphids) produced and (ii) the number of mummies from which adult wasps successfully emerged (fig 1b and 1d). Emerging wasps (1b and 1d) were also sexed in order to test whether asexual females avoid the ‘twofold’ cost of male production regardless of their mated status. To test whether asexual females that mated reproduced by facultative sex (i.e. used sperm to fertilise their eggs) I genotyped full broods of emerged wasps (fig 1b and 1d) to test for the presence of sexual alleles their offspring.

**Figure 1.**
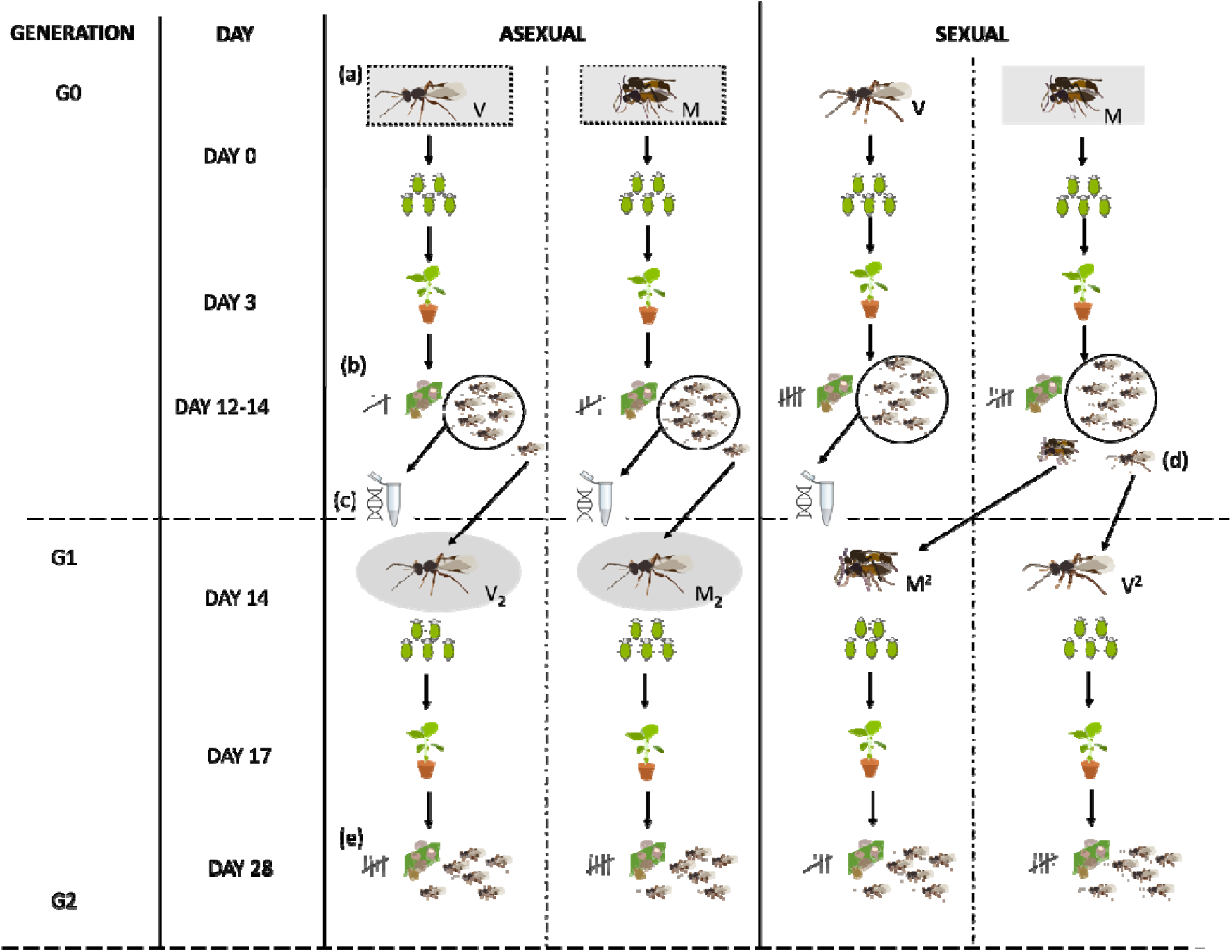
Schematic of the experimental design showing (a) the 2 treatments (virgin, V; mated, M) that sexual and asexual females were subjected to initially (G0). On day 0 mated females were paired with a male (produced by a virgin sexual female) and observed until copulation was recorded. Virgin females were kept in isolation. All wasps were then given aphid nymphs on a broad bean leaf disc to parasitise for 3 days after which the parasitised aphids were moved onto a broad bean seedling in a cellophane bag. After 14 days parasitised aphid mummies and live emerged wasps were collected (G1), (b) sexed and counted, and preserved in ethanol (c) for DNA extraction. I retained 1 live G1 asexual female per replicate from (b) to test for effects of mothers mated status on reproductive success (virgin mothers, V_2_ or mated mothers, M_2_). I took two live G1 daughters from each mated G0 sexual mother (d), one was isolated as a mummy before emergence and remained virgin (V_2_) and one was allowed to mate naturally in the cellulose bag (M_2_). These G1 females were given aphids to parasitise on the day they emerged, which, as with G0, were moved onto a broad bean seedling after 3 days. When the next generation (G2) wasps emerged (d) they, and all mummies, were counted and sexed. Boxes around focal wasps (in a and c) indicate treatments which were compared to each other to estimate the costs of male production for obligate sexuals compared to asexuals (grey rectangles) and facultative sex versus asex (grey ovals).

To assess the ‘twofold’ cost of sex (male production) and determine whether it applies to facultative (mated asexual) and obligately sexual females I looked at the number of sons and daughters produced by V, M and V_2_ and M_2_ females. To estimate this cost of sex in the case of obligate sex I compared son and daughter production between G0 obligately sexual mated M females and all (V and M) G0 asexual females (fig 1, grey boxes). To test whether the cost of male production also applies to facultatively sexual females (asexual M females) I compared son and daughter production of (i) G0 asexual V and M females (fig 1, dashed outlined boxes) and (ii) G1 asexual V_2_ and M_2_ females (fig 1 grey ovals). I predicted that the ‘twofold’ cost of sex (producing males) would only be paid by obligately sexual females (sexual M); who would produce ~50% male offspring and as a consequence would produce around half as many daughters as asexual females (asexual M and V). I predicted male production would be negligible for all asexual females, including those which mated with a sexual male (M) or whose mother had mated (M_2_). Below (in section e) I outline the statistical models applied to make these comparisons.

### (d) Experimental set up

#### Generation −1: standardised rearing

I set up a ‘G-1’ generation to standardise rearing conditions for females to be used in experiments; recently emerged females were removed from stock for all lines (7 asexual and 1 sexual) and 20 females per line were transferred onto broad bean seedlings infested with *A. fabae* in cellulose bags (Celloclair, Liestal, Switzerland). To generate enough males for the mating treatments 20 virgin sexual *Lysiphlebus* females were allowed to parasitise an aphid colony. Virgin sexual females lay only unfertilised eggs that develop as haploid males. To ensure virginity parasitised aphid ‘mummies’ were removed from infested plants in the sexual stock cage before the wasps emerged and were isolated in cellulose pill capsules. When the wasps emerged, 20 females were transferred onto broad bean seedlings infested with *A. fabae* in a cellulose bag. This process was repeated for each asexual line and the sexual population to generate focal females, but emerged females were removed from stock, and not isolated as mummies. This process was repeated with freshly emerged wasps every day for 3 days to stagger the emergence of focal wasps to increase the sample size by conducting the experiment across 3 blocks.

#### Generation 0: mating treatments

Ten to 12 days after aphids were exposed to the G-1 wasps I checked the seedlings for the presence of parasitised aphid ‘mummies’ and isolated any that I found into cellulose pill capsules. These were labelled with the line ID, whether the mother was mated or virgin (for sexuals) and the date the mummy was isolated. Mummies were checked regularly for emergence. On the peak emergence days (day 13) I selected focal females and males and assigned them to mating treatments. I set up between 3 and 11 females per treatment for each asexual line and 18 and 22 sexual females for the virgin and mated treatments respectively resulting in a total sample size of N = 126.

Asexual female wasps from 7 lines and sexual females were assigned to either mated (M) or virgin (V) treatments. For the M treatments a male produced by a sexual virgin female was introduced into the pill capsule with the female, the pair was observed continuously for 30 minutes or until copulation occurred. In *Lysiphlebus fabarum* it is possible to determine successful mating visually, the female becomes quiescent and lowers her antenna and the male is able to insert his aedeagus into her vagina. Females often reject male mating attempts in this species by curling their abdomen under and moving rapidly. For all M females I recorded whether males attempted copulation and if so whether females rejected or accepted the mating attempt. For M females that accepted copulation, the male was removed from the pill capsule and the female was retained (I did not assay the fitness of M females which did not copulate in this experiment). For the V treatments no male was added to the females pill capsule, to control for potential mechanical costs of mating, V females were gently moved around in their pill capsule using a paintbrush.

On each day that wasps were put into M and V treatments (over 3 days) once all females had been subjected to their mating treatment, they were transferred onto broad bean leaves cut into discs and embedded in 6mL 1% agarose gel in an insect rearing dish with a mesh ventilation hole (diameter, 15mm depth; SPL Life Sciences, Pocheon-si, South Korea). Leaf discs had been infested with 5 adult aphids 5 days previously. On the day of the experiments the adult aphids were removed from the leaf disc and the nymphs that they produced were counted and numbers recorded. Wasps on leaf discs were checked every day between 9:00-10:00, 12:00-13:00 and 17:00-18:00 for 3 days after they were set up and any dead females were recovered, and their time and date of death recorded. After 3 days the leaf disc was removed from the agar and transferred onto a 2-week-old broad bean seedling in a cellophane bag. If the female wasp was still alive, she was transferred onto the seedling along with the aphids.

Twelve days after the aphids had been exposed to the wasps, I cut the stem of the seedling and closed the bag (still containing the seedling) with a bulldog clip. Cutting the plant resulted in any unparasitized aphids dropping off the plant before wasps started to emerge. The bags were checked each morning from day 12 to day 16 and any emerging wasps were collected. I collected one female from each replicate (with the exception of sexual virgin females, which do not produce daughters) to start generation G1 (see below). The rest of the G1 brood were collected into 99% ethanol for genotyping. Bags containing mummies were retained and the mummies counted after all wasps had emerged.

#### Generation 1: daughters of mated and virgin females

To test whether mothers mated status influences the fitness of asexuals, I assayed the fitness of the daughters produced by G0 asexual females in the M and V treatments and G0 sexual females in the M treatment (sexual V females produced only sons). Generation G1 asexual females which had emerged into cellulose bags were collected and subjected to the same protocol as G0 females, but G1 focal females all remained virgin; they were the daughters of asexual females that had either mated (labelled M_2_) or remained virgin (labelled V_2_). I took two G1 daughters from each mated sexual G0 female, one was isolated before emergence and so remained virgin (V_2_), and the other was allowed to emerge and mate naturally in the cellulose bag (M_2_: see figure 1). Females were placed onto leaf discs for 3 days, which were then moved onto seedlings. The number of aphid nymphs and wasp survival on leaf discs was also recorded for these G1 females. I counted the number of mummies and emerged wasps produced by these G1 females. For each combination of line by treatment set up between 3 and 11 focal females, resulting in a total sample size of N = 80 G1 females.

### (d) Assaying sperm use by asexual females

To test whether asexual females used sperm from sexual males after they mated I genotyped 41 whole broods of offspring (8 broods from asexual virgin females, 29 broods from mated asexual females, and 4 broods from sexual females) using a single microsatellite marker (*Lysi07*; Sandrock et al 2007) which exhibits complete linkage with the asexuality-inducing allele (Sandrock and Vorburger 2011). DNA was extracted in a pooled reaction (up to 10 wasps from one brood per reaction) using the Qiagen DNAeasy blood & tissue kit (QIAGEN Inc, Hilden, Germany). After extraction DNA was amplified in a multiplex PCR reaction using fluorescently labelled *Lysi07* primers. PCR reactions were carried out using a Qiagen Multiplex PCR kit (QIAGEN Inc. Hilden, Germany) in 11 μl reactions and contained: 1 μL DNA template, 10 μL PCR mastermix (made up 1x QIAGEN Multiplex PCR mix and 0.1μM of each locus-specific labelled (forward) and unlabelled (reverse) primer). PCR conditions were as follows: initial denaturation at 95°C for 15 min, followed by 30 cycles of 94°C for 30 s, 58°C for 90 s, 72°C for 60 s, and final extension at 60°C for 30 min. PCR amplification was performed using a DNA Engine Tetrad ® Thermal Cycler (MJ Research, Bio-Rad, Hemel Hempstead, Herts, UK). PCR products were then sent to MRC PPU DNA sequencing and services at the University of Dundee for fragment analysis. This involves capillary electrophoresis which provides a series of different coloured peaks which vary in location on an X-axis based on the size of the microsatellite fragment. I scored these peaks using the software Geneious™. For *Lysi07*, peaks scored at 184 are found exclusively in asexual wasps and 192 in sexual wasps. When a sample had scores at both 184 and 192 it indicated the presence of both sexual and asexual alleles suggesting that an asexual wasp had mated and used sperm.

### (e) Statistical analysis

#### Parasitism failure, mummy production and wasp emergence

To test whether mated status (and reproductive mode) were related to reproductive success I ran 3 sets of generalised linear mixed models (GLMMs) with the following response variables: (i) parasitism failure (binomial, scored as 1 if no mummies were recovered; using the glmer function in the lme4 package in R; [20]), (ii) number of mummies (negative binomial, count of the number of mummies recovered from each replicate; using *glmmTMB* in R;[21]), (iii) proportion of mummies that emerged as adult wasps (binomial, modelled using the cbind function in R and using *glmmTMB*).

The G0 models included reproductive mode (sexual or asexual), mated status (virgin or mated) and their interaction effect as fixed factors. Covariates were also fitted; these were the number of aphid nymphs on the leaf disc and female wasp survival (in hours; up to a maximum of 72 hours for all wasps which survived until transfer to seedling). I also included a random effect of Line ID nested within reproductive mode. Block was not included in any model; treatments were replicated evenly across daily blocks and in no case did I find a significant effect of block on any outcome variable, nor did inclusion of block in any models influence the results. Broods that were recorded from two G0 virgin sexual females were removed from the analyses as female offspring were recovered (virgin sexual females only produce unfertilized eggs which develop as males) suggesting mislabeling.

The G1 models were similar but the data was only from asexual females, so no fixed effect of reproductive mode was fitted. The only fixed effects were mother’s mated status, (mated or virgin), survival of the focal female and number of nymphs (as covariates), and Line ID was included as a random effect. The package *DHARMa* [22] was used to check that all models were appropriately specified.

#### Daughter production and the cost of males

To compare the traditional ‘twofold’ cost of sex (male production), I tested whether obligately sexual females produced fewer daughters than all asexual females (grey boxes in fig. 1). To do this I ran a model using data from G0 females, with the number of female offspring produced as the outcome variable (negative binomial in *glmmTMB*). Data from virgin sexual females were not used for this model (as virgin sexual females produce only male offspring) but data from M sexual and asexual and V asexual G0 females was included. This G0 model included a fixed effect of reproductive mode. The number of male offspring produced was also included as a covariate, and as an interaction with reproductive mode, to test for a trade-off between son and daughter production.

I ran two additional models to test whether facultative sex (by normally asexual females) imposes a cost due to male production. These models had the same structure but included data either from only G0 or only G1 asexual females and included fixed effect of mated status (or mother’s mated status) and the number of male offspring produced (as a covariate), as well as their interaction effect. The G0 model tested whether asexual females that mated produced fewer daughters than females which remained virgin (grey boxes with dashed outlines in fig.1). The G1 model tested whether females with mated mothers produced the same number of daughters as females with virgin mothers (grey ovals in fig. 1). As with the other models mothers’ survival and number of nymphs were included as covariates and random effects were Line ID nested within reproductive mode or and Line ID only (if only data from asexual females was used).

#### Estimating overall fitness

To test whether sex, either obligate or facultative, or asexual reproduction is associated with higher reproductive success overall I used the data from G0 and G1 females (including those which failed to parasitise any aphids or produce any offspring) to estimate a standardised measure of overall fitness (*W)* for every experimental female in both generations, calculated as *PP x SP x N* where *PP* is the proportion of aphid nymphs recovered as mummies (proportion parasitised; note that if the number of mummies was higher than the number of nymphs provided, due to continued aphid reproduction after counting, this was set as 1), *SP* is the proportion of mummies that produced an adult female offspring, (successful parasitism) and *N* is the total number of offspring produced. While this method does have its shortfalls, in particular it neglects the complexities of fitness returns through males, it does provide a useful standardised way to test for differences in overall fitness according to reproductive mode. To test whether obligately sexual, facultatively sexual or asexual females have the highest fitness I ran a zero-inflated Gamma GLMM with fixed effects of ‘reproductive mode’ and ‘generation’, plus their interaction effect with ‘*W’* as the dependent variable. Reproductive mode was included as a fixed factor with 3 levels in the model, obligately sexual (M and M_2_ sexual females; V sexual females which only produce males were not included), facultatively sexual (M and M_2_ asexual females) and asexual (V and V_2_ asexual females). Generation was a fixed factor with 2 levels which indicated the mother’s generation (G0 and G1).

## 3. Results

### (a) Summary

In the first generation (G0) 103 females were offered a male to mate with (M) and 66 were not (they remained virgin, V). Of the 103 females assigned to the M treatment, 70 mated. Thirty-three M females did not mate either because males failed to initiate courtship (N = 10), or the female did not signal receptivity (N = 23). The 70 M and 66 V female wasps were given aphids to parasitise. Table 1 shows the numbers of females which remained virgin (V), mated (M), or did not mate (due to rejection or no attempt) by line.

**Table 1.** Number of first generation G0 females that remained virgin (V), accepted a mating (M: successful matings), did not accept a mating because the male did not attempt courtship, or because the female rejected the male. Also shown are the proportion of male-female introductions that resulted in a successful mating by line. A significantly lower proportion of pairings involving asexual females resulted in successful mating (*X*^2^=10.430, df = 1, p = 0.0012: asexual = 0.6, sexuals = 0.96).

| Line | N | Remained virgin | Attempted matings | Successful matings | No attempt | Rejected | proportion attempts successful |
| --- | --- | --- | --- | --- | --- | --- | --- |
| CHC17-66 | 10 | 5 | 5 | 5 | 0 | 0 | 1.00 |
| CV17-84 | 10 | 4 | 6 | 5 | 0 | 1 | 0.83 |
| IL06-658 | 8 | 3 | 5 | 5 | 0 | 0 | 1.00 |
| IL07-64 | 36 | 11 | 25 | 10 | 4 | 11 | 0.40 |
| IL09-348 | 24 | 10 | 14 | 8 | 1 | 5 | 0.57 |
| IL09-402 | 30 | 10 | 20 | 10 | 4 | 6 | 0.50 |
| IL09-554 | 10 | 5 | 5 | 5 | 0 | 0 | 1.00 |
| Sexual | 41 | 18 | 23 | 22 | 1 | 0 | 0.96 |

Of the 136 G0 females that were given aphids to parasitise 58 died before they were transferred onto the seedlings and 70 survived. In the second generation (G1), I took the daughters of 40 M and 40 V G0 asexual mothers and put them on leaf discs. Of the 80 G1 females put onto leaf discs 57 survived and 23 died before being transferred to seedlings 72 hours later (note that there was no effect of mating treatment on wasp survival in the first 72 hours in either the G0 or G1 generations: G0 (effects of reproductive mode and mated status): Log-rank *X*^2^= 1.4, df = 3, p = 0.70 and G1 (mated status only): Log-rank *X*^2^ = 2.8, df = 1, p = 0.33).

### (b) Sperm use

Forty-one broods were genotyped in total (see table S1 for a summary of brood genotypes broken down by treatment and line). All broods produced by sexual females were homozygous for the sexual allele (peak 192) at *Lysi07*. Broods produced by asexual females were significantly more likely to test positive for sexual alleles if the female had mated (M) than if she was virgin (V) (*X*^2^ =4.56, df = 1, p = 0.03); 9/30 broods produced by mated M asexual females were homozygous for the asexual allele (peak 184), 19/30 were heterozygous for sexual and asexual alleles (184/192) and the 2 were homozygous for the sexual allele (192; most likely due to a PCR failure/scoring error or misnumbering of a replicate as there is no evidence for introgression of sexual alleles in the stock lines these females were derived from). Only 1 brood produced by a virgin (V) asexual female was scored as positive (heterozygous) for the sexual allele (suggesting either mislabelling or contamination between samples); the other 7 V asexual broods were homozygous for the asexual allele.

### (c) Parasitism success

In some cases, no mummies were recovered from aphids that were exposed to *L. fabarum* females, suggesting that these females either failed to oviposit, or their eggs failed to hatch/larvae to develop resulting in complete reproductive failure. In G0 100 out of 126 females successfully parasitised aphids and produced adult offspring. A few individuals successfully produced a small number of mummies but no adult offspring (12), these females were excluded from the analyses below as I did not dissect the uneclosed mummies to determine whether the larvae died before emergence or had entered diapause. In the first generation G0, sexual females were more likely to fail to parasitise aphids than asexual females (*= 4*.*49, df = 1, 124, p = 0*.*03*). Mating had no effect on G0 females ability to parasitise and this was true for both sexual and asexual females (*= 1*.*54, df = 1,124, p = 0*.*21*. Interaction effect between reproductive mode and mated status: *= 0*.*42, df = 1, 124, p = 0*.*52*; fig. 2a).

**Figure 2.**
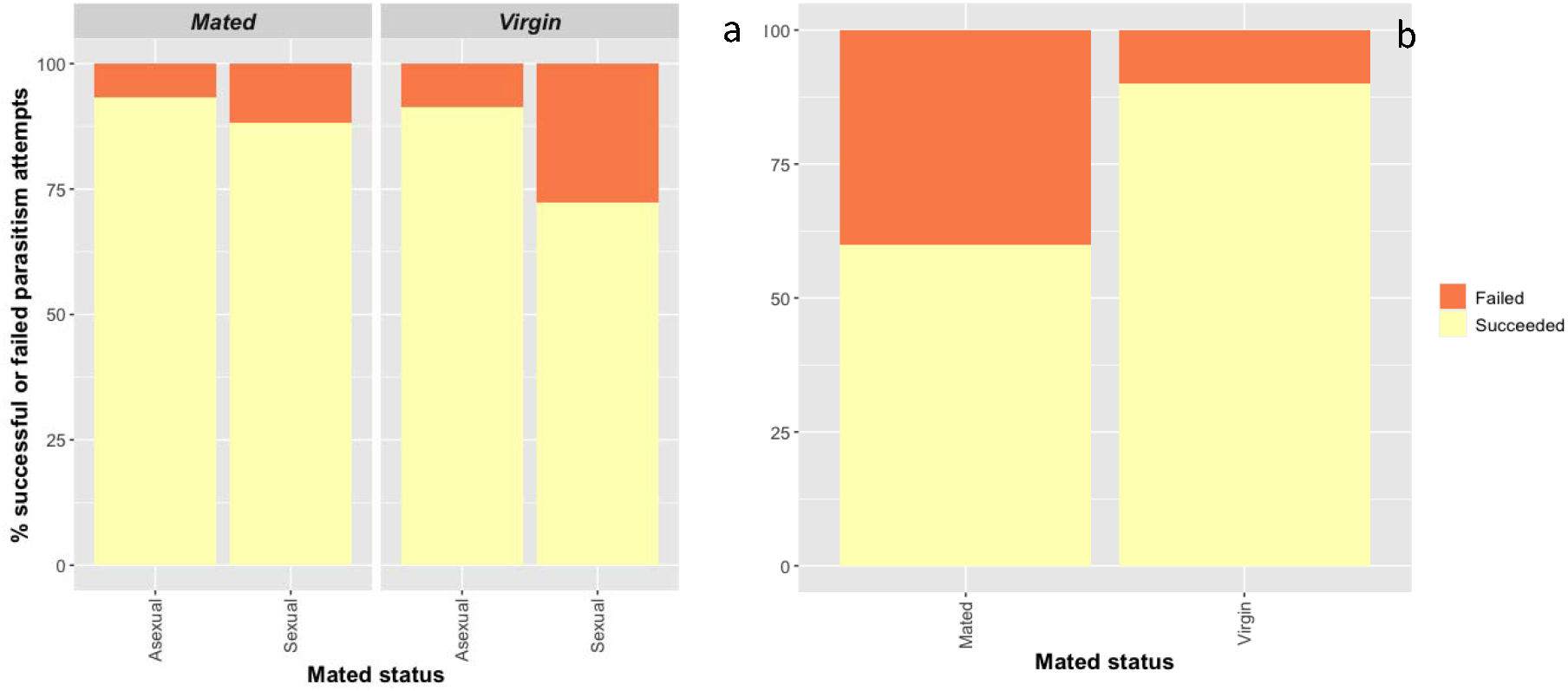
Bar plot showing the effect of mated status on % parasitism success in **(a)** G0 across 7 asexual lines and one outbred sexual line of *L. fabarum*. In G0 sexual females were more likely to fail to parasitise than asexuals but there was no effect of mated status and **(b)** G1 asexual daughters of G0 females; daughters of mated M females were significantly more likely to fail to parasitise than daughters of virgin V females.

54/80 G1 daughters of asexual G0 females successfully parasitised aphids and produced adult offspring, 6 females successfully produced a small number of mummies but no adult offspring (again, females which produced mummies but not adult offspring were removed from the analysis below). In G1 there was a large negative effect of mothers mated status on parasitism success. The G1 daughters of mated M G0 asexual females were 25% more likely to fail to parasitise aphids than the daughters of virgin V females (*= 7*.*43, df = 1,79, p = 0*.*0*.*006;* fig. 2b).

### (d) Mummy production and emergence

Although sexual females in G0 were more likely to fail to parasitise any aphids, when I considered only the females that did parasitise, sexual females were at an advantage as they produced more mummies (*= 5*.*04, df = 1, 98, p = 0*.*03;* fig. 3a) more of which produced a wasp (*= 4*.*41, df = 1, 98, p = 0*.*03;* fig. 3c) than their asexual conspecifics. Whilst sex had an effect on mummy production in G0 there was no effect of mating; virgin V and mated M females produced the same numbers of mummies (*= 0*.*30, df = 1, 98, p = 0*.*59)* regardless of whether they were sexual or asexual (interaction: *= 0*.*29, df = 1, 98, p = 0*.*59*; fig. 3a). There was an effect of mating on adult offspring production though; virgin V G0 mothers were more successful (*= 3*.*85, df = 1, 98, p = 0*.*05;* fig. 3c) for sexual and asexual mothers (interaction: *= 0*.*78, df = 1, 98, p = 0*.*38)*.

**Figure 3.**
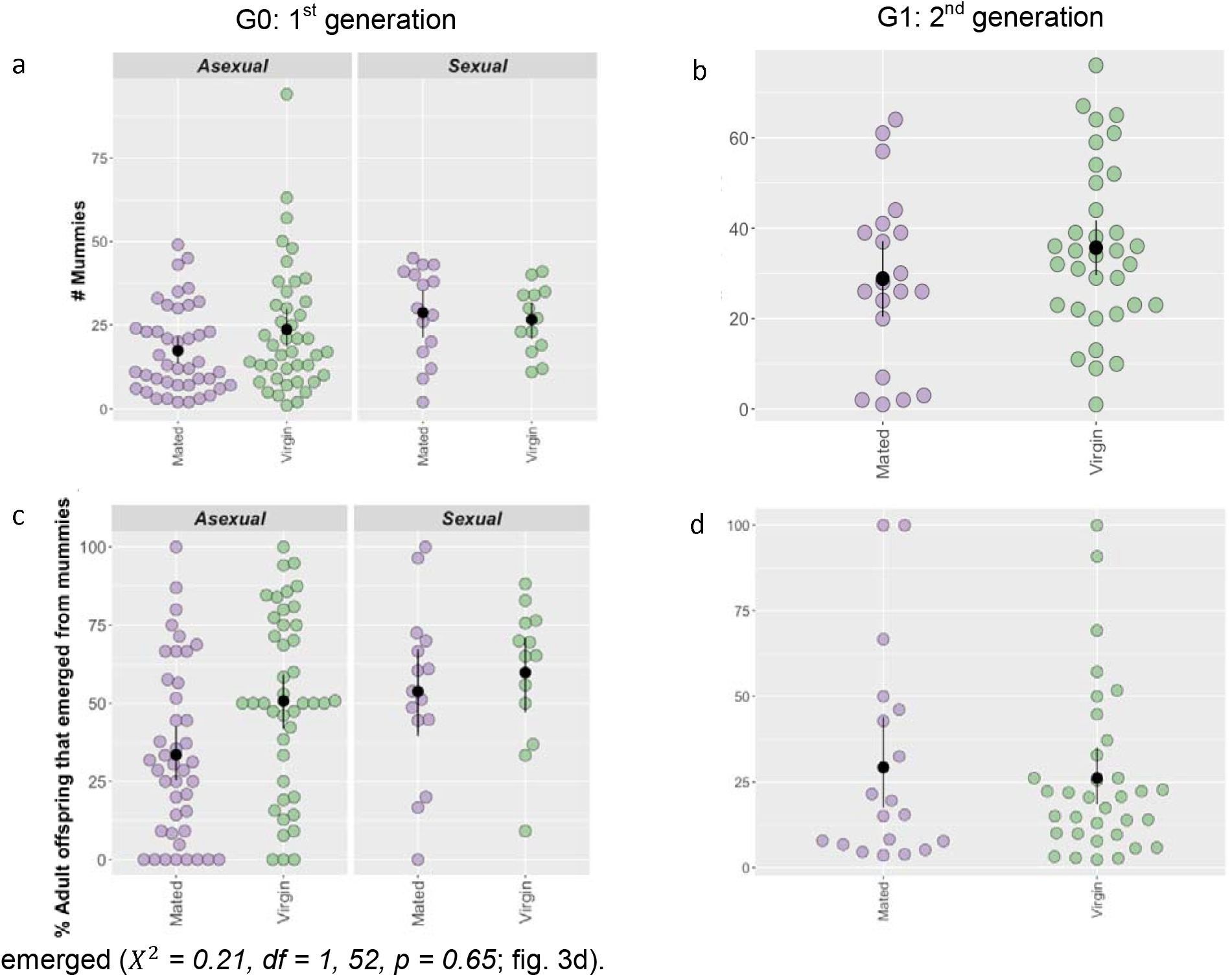
Number of mummies produced (**a & b**) and % mummies adult wasps emerged from (**c & d**) in G0 (**a & c**) and G1 (**b & d**). G0 sexual females produced more mummies than asexual females (**a**) which were more likely to emerge as adult wasps (**c**). Mummies produced by virgin G1 mothers (sexual and asexual) were also more likely to emerge as adult wasps (**c**). For asexual G1 females there was no effect of mothers mated status on mummy production (**b**) or the proportion of mummies that successfully emerged (**d**). Black points are means and error bars represent standard errors estimated by nonparametric bootstrapping.

These results were not replicated in the second generation, there was no effect of G0 mother’s mated status on the number of mummies their G1 daughters produced (*X*^2^ *= 2*.*41, df = 1, 52, p = 0*.*121*; fig. 3b) or the proportion of these mummies from which a wasp emerged (*X*^2^ *= 0*.*21, df = 1, 52, p = 0*.*65*; fig. 3d).

#### Daughter production and the cost of producing males

To test whether the classic ‘twofold’ cost of sex applies in *L. fabarum* I looked at daughter production by focal females. In G0, obligately sexual females did not produce fewer daughters than asexual females (M or V; *X*^2^ *= 0*.*02, df = 1, 98, p = 0*.*90)*. Contrary to expectations females that produced more males did not produce fewer daughters (number of sons: *X*^2^ *= 0*.*68, df = 1, 98, p = 0*.*40)*, a pattern that was the same for sexual and asexual females (reproductive mode*number male offspring: *= 3*.*23, df = 1, 98, p = 0*.*07*). I ran a second model with sexual females removed to test whether facultative sex by G0 mated asexual females imposes a cost in terms of male production, but again M and V females produced the same number of daughters (*= 0*.*49, df = 1, 71, p = 0*.*48*). Son production was very low for M and V asexual females in G0 (< 5 per mother) and there was no evidence that females which produced more males produced fewer daughters (mated status: *= 0*.*49, df = 1, 71, p = 0*.*48*; male production; *= 3*.*22, df = 1, 71, p = 0*.*07;* mated status*number male offspring *= 0*.*36, df = 1, 71, p = 0*.*55*).

I found the same pattern in the second generation (G1); male production was again very low for all asexual females and had no effect on daughter production (*= 1*.*88, df = 1, 52, p = 0*.*17*) regardless of mothers mated status (mothers mated status*number male offspring: *= 0*.*65, df = 1, 52, p = 0*.*42*). Importantly the G1 daughters of mated M G0 females (some of which are facultatively sexual) produced the same number of daughters as females produced asexually by virgin V G0 mothers (*= 2*.*24, df = 1, 52, p = 0*.*14*). Taken together these results from G0 and G1 suggest that ‘all-else-is-not-equal’ in this system, so the twofold cost of male production does not constrain the fitness of (obligately or facultatively) sexual females compared to asexuals in *L. fabarum*.

#### Estimating overall fitness through granddaughters

Fitness estimates were significantly lower for facultatively sexual females compared to obligately sexual females in G0 (z = 1.25, p = 0.04) and G1 (z = 2.37, p <0.001). Asexual and facultatively sexual females had equivalent fitness in G0 (z = 0.47, p = 0.42), the difference in fitness increased in G1, with asexual females having higher fitness, but this was not statistically different (z = 0.1.21, p = 0.059). Asexual and obligately sexual females did not differ significantly in overall fitness in G0 (z = 0.78, p = 0.27) or G1 (z = 1.17, p = 0.11; overall effect of reproductive mode: *= 16*.*23, df = 2, 90, p = 0*.*0003;* fig. 4).

**Figure 4.**
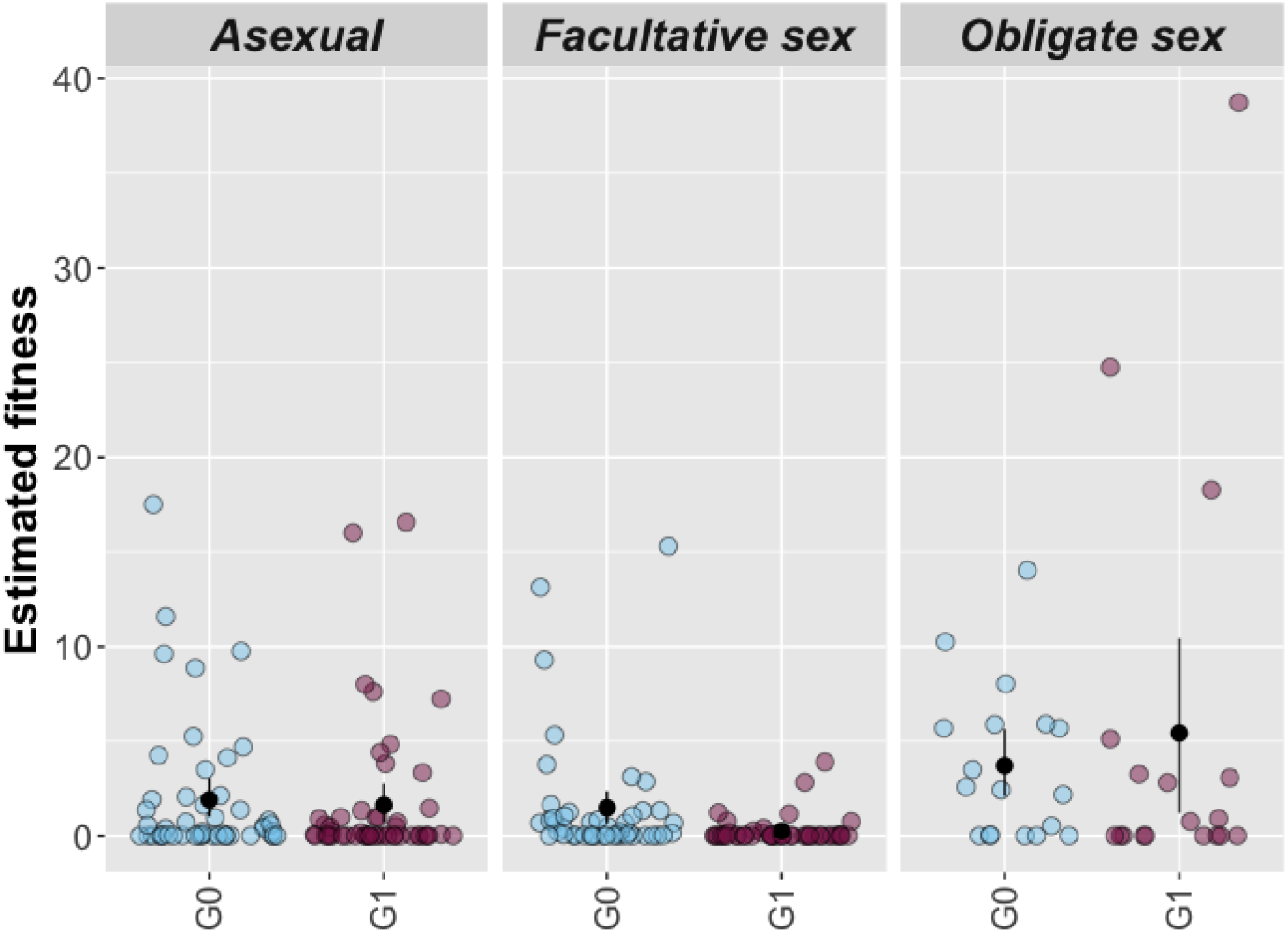
Fitness estimates for asexual (asexual V), facultatively sexual (asexual M) and obligately sexual (sexual M) G0 females. Fitness estimates were calculated by multiplying the proportion of aphid nymphs parasitized (PP), the proportion of mummies yielding adult wasps (SP), and the number of adult wasps (N). Black points represent means and error bars are standard errors estimated by nonparametric bootstrapping.

## 4. Discussion

Populations of the aphid parasitoid *Lysiphlebus fabarum* vary in their reproductive mode: asexual (parthenogenetic) reproduction dominates across Europe, but sexual populations persist in sympatry with asexuals in some regions [14,15]. In this study I show for the first time that asexual *L. fabarum* females are sexually functional and can use sperm to fertilise eggs and produce daughters. I found that between 40-100% (x□ = 60%) offemales from all 7 asexual *L. fabarum* lines assayed were facultatively sexual; they mated with sexual males and produced offspring with genetic contributions from both parents. However, asexual females that reproduced sexually did not get the best of both worlds, in terms of short-term benefits of parthenogenesis (higher daughter production) or genetic benefits of sex (increased fecundity or offspring emergence). In fact, the pattern was quite the opposite, facultatively sexual females suffered high costs of reproductive failure associated with sex *and* lower fecundity associated with asexual reproduction; they get the worst of both worlds.

The main short-term benefit of asexual reproduction is that it eliminates the ‘twofold cost of males’. In *L. fabarum*, asexuals do occasionally produce male offspring, Sandrock and Vorburger [16] found that around 1 in every 3000 asexual wasps is male in the lab. In the current study I also found low rates of male production by *all* asexual females, but crucially asexual females did not produce more daughters than sexual females rendering this cost negligible. This is because the assumption that ‘all-else-is-equal’ does not hold; obligately sexual females had higher fecundity than females from asexual lines, such that despite producing many more sons they produced the same number of daughters as asexuals, suggesting a long-term genetic benefit of sex. Females from asexual lines which engaged in facultative sex did not gain this benefit of sex though, the fecundity of asexual females was not influenced by their mated status or their mother’s mated status. There was, however, a significant fitness cost to sex, it increased the risk of reproductive failure by 12% compared to parthenogenesis. This cost was even higher for asexual females that mated and reproduced sexually; when asexual G0 females mated and could reproduce sexually, 40% of their G1 daughters failed to reproduce compared to only 10% of the G1 daughters of asexual G0 females that did not mate. When all measures of reproductive success were combined to estimate a standardised measure of fitness asexual and obligately sexual females did not differ, which may explain why sexual and asexual populations exist in nature. Facultatively sexual females on the other hand had lower fitness, presumably due to the combined effects of high reproductive failure associated with sex, and lower fecundity associated with parthenogenesis (asex).

Asexual reproduction is thought to have first evolved approximately 0.5 million years ago in *L. fabarum*, arising through the inheritance of a single recessive allele which results in the haploid products of meiosis I fusing to form diploid gametes which develop without fertilisation (central-fusion automixis; [17]). Although asexuality may have arisen 0.5 million years ago it is unlikely that the asexual lineages I tested have existed for so long without sex and recombination (see [17]). The asexual lines used in this experiment have, however been reared in the lab for 7-18 years without sex. Despite the lack of sex (potentially for up to 400 generations) some females from all seven asexual *L. fabarum* populations exhibited sexual behaviour and were able to integrate paternal DNA into the female offspring they produced. The maintenance of sexual traits in *L. fabarum* makes them unique compared to many other asexual eukaryotes, where sexual traits such as mate location, attraction, allowing copulation and storing sperm decay rapidly due to high costs of maintenance [23-26].

It is unclear is why sexual traits have persisted in asexual *L. fabarum*as I found that facultative sex is costly for reproductive success in this species; 40% of the daughters of mated asexual females failed to reproduce entirely, which should result in strong selection on asexual females that do not invest in sexual signalling or accept matings. There are multiple possible reasons that sex, in particular facultative sex, might increase the risk of reproductive failure. One explanation for is that some of the daughters of some daughters of G0 M asexual females are triploid with low fertility. Triploid females may represent the proportion of the females in G1 that failed to produce any offspring. Whilst I was unable to detect triploids using multiplex PCRs here (as broods were pooled) Sandrock et al [15] found that around 2% of females in all-female broods were triploid in asexual *L. fabarum* in the field, supporting this possibility. However, triploidy is not a given under central-fusion automixis as eggs are laid when they are in metaphase I, but diploidy is not restored until anaphase II [17]. Another non-mutually exclusive possibility is that ‘genetic slippage’ occurs, resulting in lower fitness of sexually produced offspring. Genetic slippage, essentially outbreeding depression, occurs when combinations of genes that normally work well together are broken up by sex, creating new genotypes which have low fitness [3–8,12]. Sex and recombination can produce novel multilocus genotypes that perform better than the parental genotypes in some environments, but genetic slippage can mean that they perform much worse [12]. In this study it may be that the new genotypes created by sex were costly for *L. fabarum*, and this cost may have been more severe under facultative sex because of the greater genetic differentiation between asexual females and sexual males. Follow up studies which replicate this experiment, but genotype individual mothers and offspring are needed to shed light on the extent to which triploidy leads to reproductive failure. Similarly, repeating this study but using rare males that are produced by asexual females will allow for more explicit tests of the genetic slippage hypothesis, as facultative sex with more closely related mates should in theory be less costly.

The mating systems and population dynamics characteristic of aphid parasitoids like *Lysiphlebus* might explain why facultative sex has persisted in *L. fabarum* despite the costs. Parasitoid abundance increases over the season as hosts become more numerous, until populations crash later in the summer [27]. At the end of the season the abundance of *L. fabarum* females in parasitised aphid colonies can become very high which could have a number of effects on the rate of facultative sex and its consequences. First, high population densities are expected to increase the likelihood that normally asexual females will encounter and mate with rare asexual males (or sexual males in regions of sympatry).

Second, mating between kin is common in parasitoids because many females parasitise aphids in the same colony that they emerged from (they are quasigregarious), which may counter the detrimental effects of genetic slippage (i.e. they are less likely to produce low fitness recombinants with closely related mating partners: [28]). Genetic slippage is essentially a form of severe outbreeding depression [29], and so its costs may be reduced under inbreeding. Third, the main cost of sex (increased reproductive failure), may be relaxed at certain times in the season. For instance, during the ‘mid-season crash’ aphid populations become locally extinct over just a few days, in part due to high natural enemy pressure [30]. At such times the cost of sex may be less significant if the costs of host-finding and host competition constrain parasitoid fitness to a greater extent.

In addition to relaxed costs, facultative sex could also be an important ‘bet-hedging’ strategy for *L. fabarum* late in the season if it reduces the variation in fitness across environments compared to parthenogenesis. Bet-hedging has been proposed to maintain traits that are costly for individual fitness, like sex (and polyandry [31,32]) ensuring the genes underlying these traits persist even if the traits themselves can be costly to the bearers. Williams [33] used the analogy of reproduction as a lottery to explain sex as bet-hedging: in this analogy asexual reproduction is like buying many lottery tickets with the same number, if you win, you win big. Reproducing sexually, you don’t buy so many tickets, but they have different numbers, increasing the chances you win, but reducing the amount you win. Sex and recombination result in more genetically diverse offspring, which differ from each other and their parents. By producing sexual offspring with more diverse genotypes, some of them will have low fitness in the same conditions as their parents. The genotypes that perform poorly in the parental environment may be better adapted to different conditions, they may thrive when the environment changes [8,33]. Asexual reproduction on the other hand tends to result in offspring that are genetically very similar, if not identical, to their parents, and so do very well in the same conditions their parents experienced, but if the environment changes, they are more likely to fail to reproduce entirely. Under bet-hedging, sex is a risk-averse strategy associated with moderate fitness under a range of environments, while asexual reproduction could provide big benefits under a stable environment but is risky if conditions change. The scenario that aphid parasitoids like *L. fabarum* face at the end of the season may increase the importance of facultative sex as a bet-hedging strategy because some offspring will need to survive overwinter as diapause pre-pupae and reproduce the following spring, in drastically different environmental conditions from their parents.

When viewed in the light of the natural history of aphid parasitoids like *L. fabarum* the results of this study suggest that relatively frequent sex, potentially between closely related males and parthenogenetic females, may be responsible for maintaining high levels of genetic variation and heterozygosity in asexual *L. fabarum*. If sex occurs mainly late in the season it is also likely to be fairly synchronous, mirroring the reproductive phenology of their cyclically parthenogenetic aphid hosts. When normally parthenogenetic individuals all reproduce sexually at the same time in this way, Kokko [11] has shown that facultative sex is more stable and robust to the invasion of obligate asexual reproduction. Synchronous seasonal sex between siblings may explain not only the how heterozygosity and allelic diversity is maintained in asexual *L. fabarum* [14,15] but also why sex and its associated traits have persisted. Studies that measure the frequency of sex in the field, and its associated costs will reveal the full extent of facultative parthenogenesis and its evolutionary significance in *L. fabarum*, so that we can establish whether facultative sex has in this case made asexual reproduction more successful.

## Supporting information

Table S1 Sexual alleles in asexual broods

## Data, script and code availability

All data, code and supplementary information can be accessed here: https://osf.io/qb7yz/?view_only=bbb5677d9ce14c6dab3ac42b94345c17

