## Supplementary material for "Is facultative sex is the best of both worlds in the parasitoid wasp *Lysiphlebus fabarum?*": Table S1 Sexual alleles in asexual broods

**Table S1** Broods produced mated M females across all lines that were genotyped at *Lysi07* and for which the sexual allele (192) was scored as present indicating sperm was used (for asexual broods).

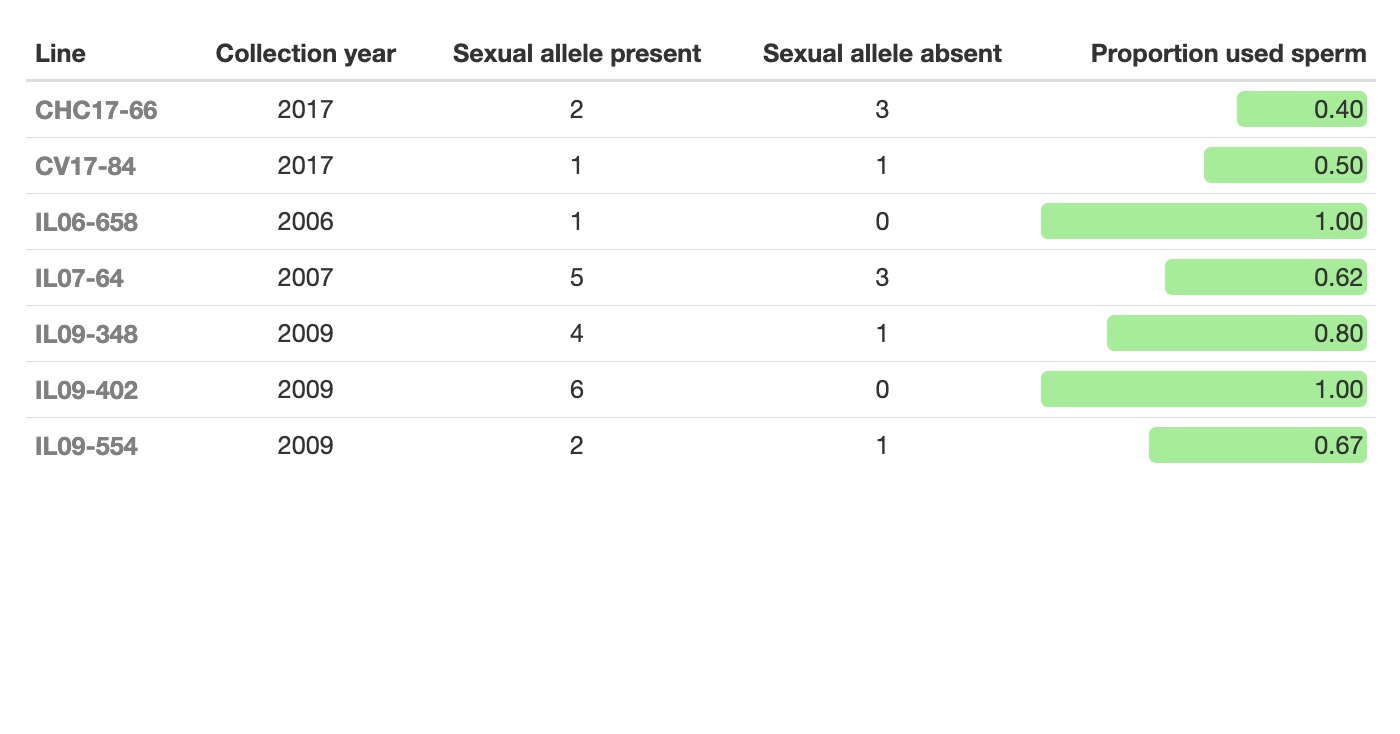
